## Supplementary Text S1 for "FiCoS: a fine-grained and coarse-grained GPU-powered deterministic simulator for biochemical networks"

#### 1 Runge-Kutta methods

In this section we provide the mathematical notation at the basis of numerical integration methods for systems of Ordinary Differential Equations (ODEs). Given a system of $n$ ODEs, its Cauchy problem is defined as:

$$\begin{cases} \frac{d\mathbf{X}}{dt} &= f(t, \mathbf{X}) \\ \mathbf{X}(t_0) &= \mathbf{x}_0, \end{cases} \tag{1}$$

where $\mathbf{X}(t) = (X_1(t), X_2(t), \ldots, X_N(t))$ and $t \in [t_0, t_{max}]$.
A generic Runge-Kutta method of order $s$ and stages $r$ is defined as:

$$\mathbf{X}(t_{n+1}) \simeq \mathbf{x}_{n+1} = \mathbf{x}_n + h \sum_{i=1}^{s} \beta_i k_i, \tag{2}$$

where $h$ is the integration step-size used during the resolution of the ODE system. The auxiliary variables $k_i$ are given by the following relationship:

$$k_i = f(t_n + c_i h, \mathbf{x}_n + h \sum_{j=1}^{s-1} \alpha_{ij} k_j), \text{with } i = 1, \ldots, s. \tag{3}$$

The coefficients $\alpha_{ij}$, $\beta_i$ and $c_i$ allow for characterizing every Runge-Kutta method, which can be represented in the so-called Butcher tableau (see Table 1).

Table 1: Butcher tableau of a generic Runge-Kutta method

$$\begin{array}{c|cccc} c_1 & \alpha_{11} & \alpha_{12} & \cdots & \alpha_{1s} \\ c_2 & \alpha_{21} & \alpha_{22} & \cdots & \alpha_{2s} \\ \vdots & \vdots & \vdots & \ddots & \vdots \\ c_s & \alpha_{s1} & \alpha_{s2} & \cdots & \alpha_{ss} \\ \hline & \beta_1 & \beta_2 & \cdots & \beta_s \\ & \beta_1^* & \beta_2^* & \cdots & \beta_s^* \end{array} = \begin{array}{c|c} \mathbf{c} & \mathbf{\Lambda} \\ \hline & \boldsymbol{\beta}^T \end{array}$$

The Runge-Kutta methods are partitioned in three classes based on the coefficients represented in the Butcher tableau. To be more precise, the following rules are applied to classify every Runge-Kutta method:

- if the matrix $\mathbf{\Lambda}$ is lower triangular (i.e., $\alpha_{ij} = 0$ for $j > i$, with $i, j = 1, \ldots, s$), then the method is said to be *explicit*;
- if the matrix $\mathbf{\Lambda}$ is lower triangular, including the main diagonal, the method is called *semi−implicit*;

Table 2: Butcher tableau containing the DOPRI5 coefficients (left) and RADAU5 coefficients (right)

| | | | | | | | |
|---|---|---|---|---|---|---|---|
| $0$ | | | | | | | |
| $\frac{1}{5}$ | $\frac{1}{5}$ | | | | | | |
| $\frac{3}{10}$ | $\frac{3}{40}$ | $\frac{9}{40}$ | | | | | |
| $\frac{4}{5}$ | $\frac{44}{45}$ | $\frac{-56}{15}$ | $\frac{32}{9}$ | | | | |
| $\frac{8}{9}$ | $\frac{19372}{6561}$ | $\frac{-25360}{2187}$ | $\frac{64448}{6561}$ | $\frac{-212}{729}$ | | | |
| $1$ | $\frac{9017}{3168}$ | $\frac{-355}{33}$ | $\frac{46732}{5247}$ | $\frac{49}{176}$ | $\frac{-5103}{18656}$ | | |
| $1$ | $\frac{35}{834}$ | $0$ | $\frac{500}{1113}$ | $\frac{125}{192}$ | $\frac{-2187}{6784}$ | $\frac{11}{84}$ | |
| | $\frac{71}{57600}$ | $0$ | $\frac{71}{16695}$ | $\frac{71}{1920}$ | $\frac{-17253}{339200}$ | $\frac{22}{525}$ | $\frac{-1}{40}$ |
| | $\frac{-12715105075}{11282082432}$ | $0$ | $\frac{87487479700}{32700410799}$ | $\frac{-10690763975}{1880347072}$ | $\frac{701980252875}{199316789632}$ | $\frac{-1453857185}{822651844}$ | $\frac{69997945}{29380423}$ |

| | | | |
|---|---|---|---|
| $\frac{4-\sqrt{6}}{10}$ | $\frac{88-7\sqrt{6}}{360}$ | $\frac{296-169\sqrt{6}}{1800}$ | $\frac{-2+3\sqrt{6}}{225}$ |
| $\frac{4+\sqrt{6}}{10}$ | $\frac{296+169\sqrt{6}}{1800}$ | $\frac{88+7\sqrt{6}}{360}$ | $\frac{-2-3\sqrt{6}}{225}$ |
| $1$ | $\frac{16-\sqrt{6}}{36}$ | $\frac{16+\sqrt{6}}{36}$ | $\frac{1}{9}$ |
| | $\beta_1$ | $\beta_2$ | $\beta_3$ |
| | $\beta_1^*$ | $\beta_2^*$ | $\beta_3^*$ |

- conversely the method is *implicit*. In this case, in order to calculate the auxiliary variables $k_i$, with $i = 1, \dots, s$, at least one non-linear system must be solved. As a matter of fact, each $k_i$ depends on all $k_j$, with $j = 1, \dots, s$. Generally, implicit Runge-Kutta methods are exploited for the resolution of stiff problems due to their stability regions [1].

In contrast to the multi-step methods, Runge-Kutta methods are single-step methods in which it is possible to change the step-size during the ODE resolution, according to the desired error, controlled by means of two tolerances (i.e., absolute and relative). The absolute tolerance $\varepsilon_a$ is a threshold below which the value of the solution is considered unimportant, determining the accuracy when the solution approaches zero. The relative tolerance $\varepsilon_r$ is a measure of the error relative to the size of the solution. Approximately, this value controls the number of correct digits in the solution, except those smaller than $\varepsilon_a$. Adaptive Runge-Kutta methods are characterized by two embedded methods, as shown in the Butcher tableau (Table 1) with $\beta_i$ and $\beta_i^*$, with $i = 1, \dots, s$.

FiCoS is based on two different integration methods belonging to the Runge-Kutta family: the DOPRI method [2, 3, 4] exploited in the absence of stiffness, and the Radau IIA method [5, 6] when the system is stiff. To be more precise, we exploited DOPRI5 and RADAU5, which are explicit and implicit adaptive Runge-Kutta methods of order 5, respectively, capable of varying the integration step-size during the resolution of the ODE system. In Table 2 are depicted the coefficients of the embedded methods characterizing DOPRI5 (left) and RADAU5 (right). For DOPRI5, we exploited the coefficients suggested in [7], which were specifically proposed to design a DOPRI5 variant implementing the dense output feature, also known as continuous extension [8]. Notice that also the RADAU5 variant implemented in FiCoS uses a dense output solution. The dense output feature was exploited to avoid the need to stop the integration methods in correspondence to the simulation time instants specified by the user. These output time instants are calculated by means of interpolation functions, such as cubic Hermite spline, without any other additional function evaluations.

In order to reduce the possible round-off errors, the Butcher tableau of RADAU5 is modified following a very simple idea (see [5] for additional details), obtaining the following coefficients for the embedded methods: $\beta_1 = 0$; $\beta_2 = 0$; $\beta_3 = 1$; and $\beta_1^* = -13 - 7\sqrt{6}$; $\beta_2^* = -13 + 7\sqrt{6}$; $\beta_1^* = -1$.

RADAU5 required the resolution of one non-linear system for each $k_i$ (with $i = 1, \dots, s$) that must be calculated. All these non-linear system are converted into linear systems exploiting the Newton—Raphson method [9], and then solved by means of the LU factorization method [10]. Notice that in each iteration of the Newton—Raphson method, these linear systems must be solved.

In both the algorithms, once the solutions of the two embedded methods are calculated, they are used to evaluate the committed error. If the error is greater than the desirable error—which depends on the user-defined absolute and relative tolerances—the current solutions are rejected and re-calculated using a smaller step-size, otherwise they are accepted. In both cases, the solutions of the two embedded methods are also used to calculate the step-size for the next iteration.

### 2 Compute Unified Device Architecture

In order to develop FiCoS, we relied on the NVIDIA Compute Unified Device Architecture (CUDA), which represents the most widespread GPU computing library. CUDA was proposed by NVIDIA to

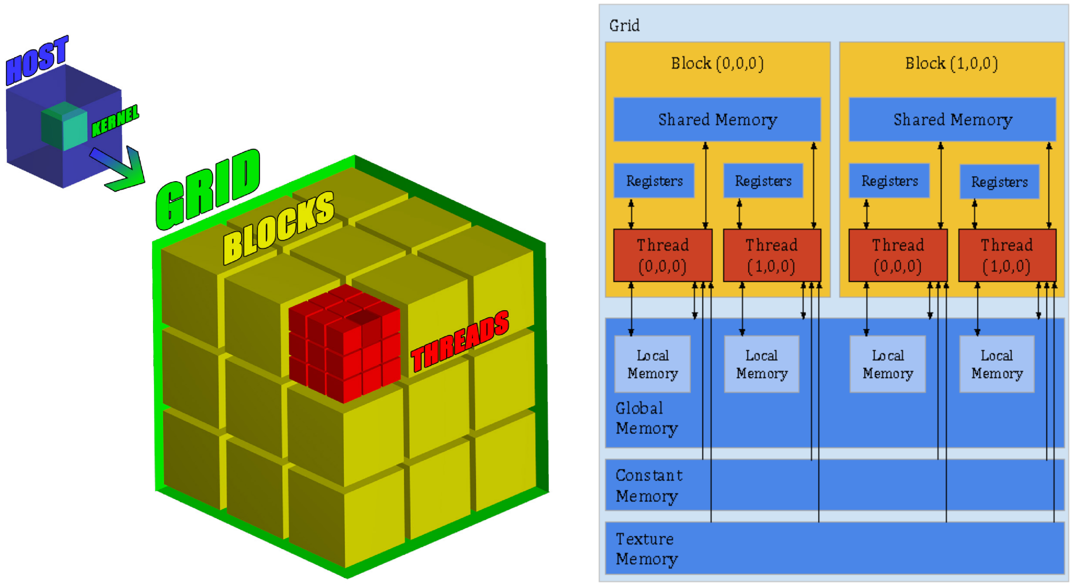


Figure 1: Execution and memory hierarchies of the CUDA's architecture *Left panel* depicts the execution hierarchy, where a single kernel is launched by the CPU (host) and run on the GPU (device) by using multiple threads. When the host launches a kernel, a three-dimensional structure, named grid (green cube) is generated by the device. This grid is partitioned in three-dimensional structures called blocks (yellow cubes), which contain the threads (red cubes). Notice that the number of threads depends on the dimensions of blocks and grids, which must be provided by the programmer. As soon as the grid is created, the device automatically schedules each block on one of the free streaming multiprocessor that are available. Since the GPUs have a different number of streaming multiprocessor, this mechanism allow for a transparent scaling of performances. *Right side* shows the memory hierarchy, which is composed of many different memories with different scopes. Registers and local memories represent the only private memories in which every thread saves its own data. The low access latency shared memory can be used by the threads belonging to the same block to communicate and share information/data. The global memory can be accessed by all the threads as well as by the host, so it is used to the CPU-GPU communications (and vice-versa). Notice that this memory is the largest one, but also the slowest due to its high latency, which was mitigated by means of a cache starting from the Fermi architecture. Texture and constant memories can be assessed only in read by the threads while the host can write the data that are kept constant during the execution of the kernel.

exploit their GPUs for general-purpose computational tasks, following the General-Purpose Computing on Graphics Processing Units (GPGPU) paradigm. Despite the availability of several libraries and high-level Application Programming Interfaces (APIs) provided by CUDA, the direct porting of the CPU code is unfeasible and represents the main challenge of GPGPU computing. As a matter of fact, programmers have to redesign and rewrite the code to fully exploit of the computational power and the massive parallelism of GPUs. In order to develop GPU code, programmers write different kernels (`C/C++`) that are launched by the CPU (called host) and run on the GPU (called device). Following an execution hierarchy, once a kernel is loaded, the device automatically generates a three-dimensional structures named grid that is divided into different three-dimensional structures called blocks containing the threads, which represent different copies of the kernel (see Fig. 1, left panel). The possible conditional divergences among the threads are managed by CUDA by combining the Single Instruction Multiple Data (SIMD) architecture along with a flexible multi-threading strategy. Besides the execution hierarchy described above, CUDA is based on a memory hierarchy as schematically represented in the right panel of Fig. 1. The memories are partitioned as follows: ($i$) the global memory, which is accessible from all threads, is the largest one (a few GBs), but at the same time is the slowest one even if the introduction of a cache level starting from the Fermi architecture; ($ii$) the shared memory is much smaller, up to 112 KB for each streaming multi-processor and limited to 48 KB for each block running over the streaming multiprocessor, but extremely faster with respect the global memory. This memory is used to communicate and share information/data among the threads belonging to the same block; (*iii*) the constant memory, which is cached and read only, is very small (i.e, up to 10 KB for each multi-processor) but can be exploited to store the data that are kept constant during the execution of the kernel; (*iv*) the local memories are represented by private registers and arrays in which the thread saves its own data.

Given these characteristics of the CUDA architecture, the computational power of these many-core devices can be obtained only by optimizing both the thread partitioning and memory usage along with the redesign of the algorithms in a set of appropriate kernels.

### 3 GPU implementation of FiCoS

In this section, we provide a detailed description of the GPU implementation of FiCoS.

#### 3.1 Data structures and CUDA memory usage

As a first step, starting from the RBM given as input, FiCoS automatically generates the systems of ODEs and encodes both the matrices $\mathbf{H} = (\mathbf{B} - \mathbf{A})^T$ and $\mathbf{A}$ as two arrays of `short2` CUDA vector type, named $\mathbf{H_v}$ and $\mathbf{A_v}$, respectively. CUDA vector types are multi-dimensional arrays with different components, ranging from 1 to 4, which are addressed by `.x`, `.y`, `.z`, and `.w` components, respectively. Note that we exploited the `short2` CUDA vector type because it is 4-aligned in the memory, meaning that a single instruction in required to fetch a whole entry. We used these data structures to compress and store the matrices $\mathbf{A}$ and $\mathbf{H}$, which are highly sparse, by removing all zero elements to save memory and at the same time to avoid unnecessary readings from the global memory of the GPU. The following strategy is applied to generate the CUDA vector type $\mathbf{H_v}$ and $\mathbf{A_v}$:

- for each non-zero element $h_{ji}$ of $\mathbf{H}$, with $i = 1, \ldots, M$ and $j = 1, \ldots, N$, we store in the `.x` and `.y` components of $\mathbf{H_v}$ the values $i$ and $h_{ji}$, respectively;
- for each non-zero element $a_{ij}$ of $\mathbf{A}$, with $i = 1, \ldots, M$ and $j = 1, \ldots, N$, we store in the `.x` and `.y` components of $\mathbf{A_v}$ the values $j$ and $a_{ij}$, respectively.

Besides these two data structures, we use two additional arrays of `int` type, named $\mathbf{O_H}$ and $\mathbf{O_A}$, to parse $\mathbf{H_v}$ and $\mathbf{A_v}$ inside the GPU. $\mathbf{O_H}$ and $\mathbf{O_A}$ store the offsets to correctly fetch the entries of $\mathbf{H_v}$ and $\mathbf{A_v}$, respectively. Each thread $j$, with $j = 0, \ldots, N - 1$, reads the $j$ and $j + 1$ elements of $\mathbf{O_H}$, whose values indicate the first index and the last index (minus 1) to access the $\mathbf{H_v}$ structure. Similarly, $\mathbf{O_A}$ stores the indexes that allow the threads to correctly decode the $\mathbf{A_v}$ structure. Finally, an array $\mathbf{K}$ of type `double` is used to store the values of the kinetic constants. An example of these data structures and their decoding is depicted in Fig. 2.

These CUDA structures allow for improving the performance of FiCoS at two different levels. First, a single instruction is sufficient to either load or store a multi-word vector. This means that the total instruction latency for a particular memory transaction is lower, while the bytes per instruction ratio is higher. Second, by using vector types, a transfer request from a warp has a larger net memory throughput per transaction, yielding a higher bytes per transaction ratio. Therefore, with a lower number of transfer requests, the memory controller can reduce the contentions and achieve a higher overall memory bandwidth utilization.

#### 3.2 Execution workflow

Once the data structures encoding the system of ODEs are generated from the input files (see Supplementary Text S2), FiCoS solves the systems of ODEs by means of the DOPRI5 method [2, 3, 4] in the absence of stiffness or the RADAU5 method [5, 6] when the system is stiff. We point out that our implementation was inspired by the source code of Blake Ashby who ported the original `Fortran` code of Hairer and Wanner [5, 4] to `C++`.

Starting from an initial time instant $t_0$, the system of ODEs is integrated up to a given maximum simulation time $t_{max}$. The dynamics of the species appearing in the RBM, described by the ODEs, are sampled and saved at specified time instants within the interval $[t_0, t_{max}]$.

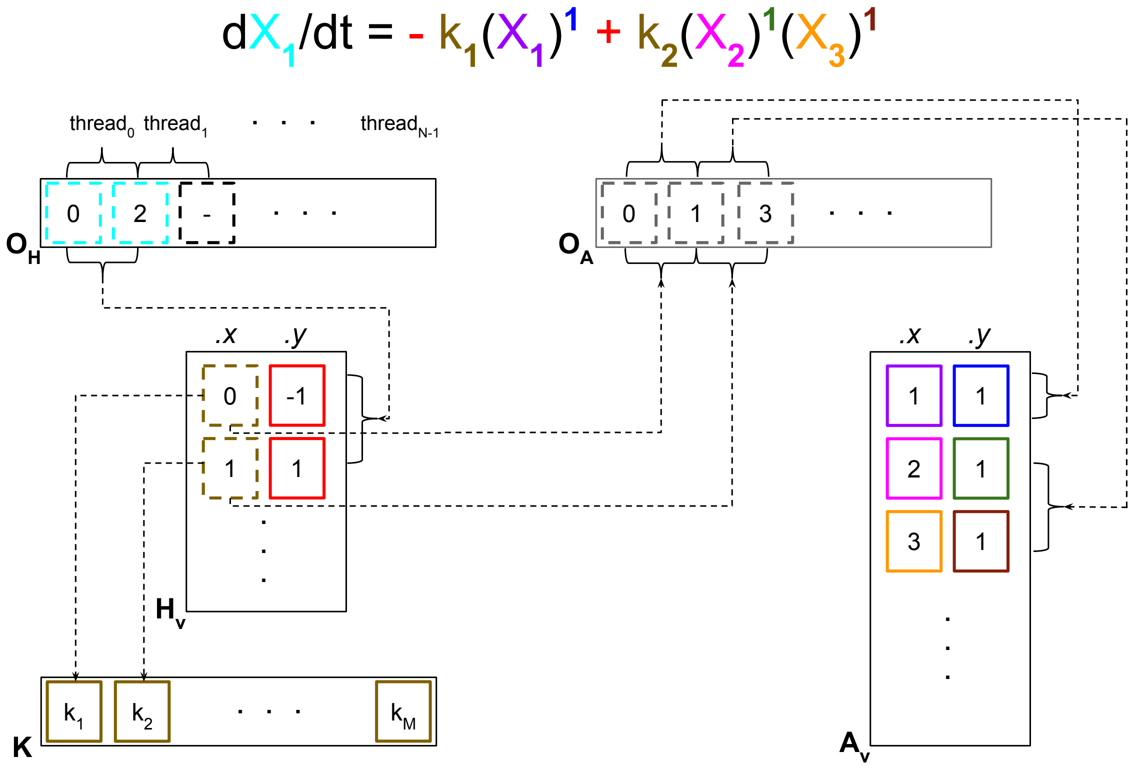


Figure 2: The monomials composing the polynomial function that describes the ODE of the species $X_1$, at the top of the figure, are encoded in $\mathbf{O_H}$, $\mathbf{O_A}$, $\mathbf{H_v}$, $\mathbf{A_v}$ and $\mathbf{K}$. A detailed description of the procedure to decode the data structures is provided below. Note that the ODE is automatically generated using the data structure components with solid borders; the colors used to highlight the terms composing the ODE are also used in the data structure components. Each thread $j$, with $j = 0, \dots, N-1$, reads the $j$ and $j+1$ elements of $\mathbf{O_H}$ (denoted by the lightblue borders). Each thread $j$ fetches the values stored in $\mathbf{H_v}$, starting from the row indicated by the value of the element $j$ of $\mathbf{O_H}$ up to the row indicated by the value (minus 1) of the element $j+1$ of $\mathbf{O_H}$. In this example, thread 0 reads the `.x` and `.y` components of the first two rows (i.e., rows 0 and 1) of $\mathbf{H_v}$. The `.x` component of each row of $\mathbf{H_v}$ (denoted by dark yellow) indicates both the position of the vector $\mathbf{K}$—which contains the values of the kinetic constants—and the first index of $\mathbf{O_A}$ where each thread $j$ must read; the `.y` component (red borders) codifies both the sign and coefficient of the monomial. In this example, the `.x` and `.y` components of $\mathbf{H_v}$ codify the coefficients $-1k_1$ and $+1k_2$ of the first and the second term of the ODE, respectively. Afterwards, each thread $j$ fetches the values encoded in $\mathbf{O_A}$ starting from the value stored in the position indicated by the index previously read up to value (minus 1) stored in the next position. The values stored in the `.x` (purple, magenta, and orange borders) components of $\mathbf{A_v}$ are the indexes of the species, while the `.y` (blue, dark green, and dark red borders) components codify the stoichiometric coefficients. In this example, the row 0 of $\mathbf{A_v}$ contains the factor $(X_1)^1$, while the rows 1 and 2 allow the thread 0 to generate the factor $(X_2)^1(X_3)^1$. By so doing, the thread 0 is capable of reproducing the polynomial composing the ODE: $-k_1(X_1)^1 + k_2(X_2)^1(X_3)^1$.

FiCoS's workflow can be summarized in 5 distinct phases, as depicted in Fig. 3. Note that only phases $P_1$ and $P_5$ are executed by the host (light orange boxes in Fig. 3), while the others are executed by the device (light green boxes in Fig. 3). Phases $P_2$, $P_3$ and $P_4$ correspond to three different kernels ($K_1$, $K_2$, and $K_3$) developed to take advantage of the coarse-grained parallelization strategy. Moreover, these phases recall other lightweight kernels, developed to fully leverage the parallel architecture of the modern GPU, which exploit the dynamic parallelism (DP) [11] to realize the fine-grained parallelization strategy of the aforementioned numerical integration methods. We describe hereafter each phase with its CUDA kernels to explain our novel parallelization strategy that allows FiCoS to achieve the relevant speed-up shown in the Results section.

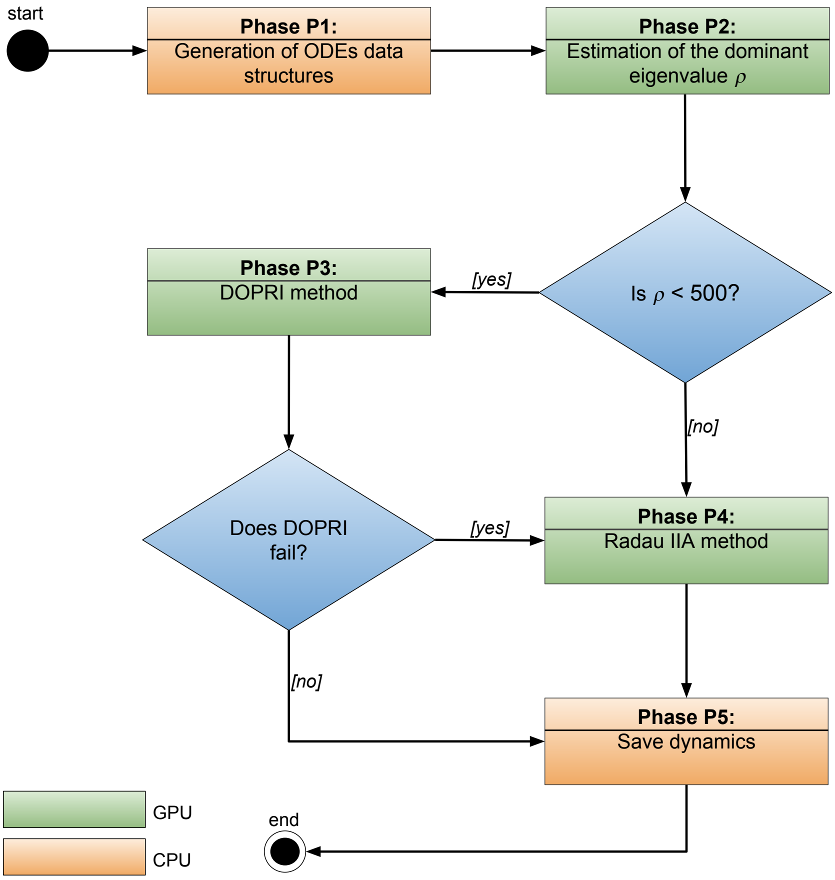


Figure 3: Simplified scheme of FiCoS workflow. During phase $P_1$ all the data structures used to encode the system of ODEs are generated. In phase $P_2$ each thread $\phi$, with $\phi = 0, \ldots \Phi - 1$ estimates the dominant eigenvalue $\rho$ related to the simulation $\phi$, using the the $\phi$-th parameterization and the $\phi$-th set of initial concentrations. All threads whose dominant eigenvalue $\rho$ is less than 500 are added in the set $\mathcal{L}_{DOPRI}$, which contains the threads that use DOPRI as integration algorithm (phase $P_3$). If some simulations using DOPRI fail, those threads are added to the set $\mathcal{L}_{RADAU}$, which contains the threads that use Radau IIA as integration algorithm (phase $P_4$). As soon as the phases $P_3$ and $P_4$ are completed, the output data (i.e., the concentration values of molecular species sampled at fixed time instants) are transferred to the host that writes them in output files (phase $P_5$).

**Phase $\mathbf{P_1}$.** It is executed by the host and implements the generation of the data structures encoding the system of ODEs. For further details, see the Data structures and CUDA memory usage Section.

**Phase $\mathbf{P_2}$.** This phase implements the estimation of the dominant eigenvalue $\rho$ of the Jacobian matrix $\mathbf{J}$ by means of the norm bound of $\mathbf{J}$ itself. As norm bound, we used the so-called maximum absolute row sum norm, defined as:

$$||\mathbf{J}||_\infty = \max_{1 \leq i \leq N} \sum_{j=1}^{N} |J_{ij}|. \tag{4}$$

Note that the dominant eigenvalue $\rho$ is equal to the spectral radius, which is defined as the largest absolute value of the eigenvalues of $\mathbf{J}$.

Since the value $\rho$ strictly depends on both the initial concentrations and the kinetic parameter values, $\rho$ is calculated for each simulation $\phi$, with $\phi = 0, \ldots \Phi - 1$, which must be performed. This phase is implemented using 2 different CUDA kernels:

- kernel $K_1$: each thread $\phi$, with $\phi = 0, \ldots \Phi - 1$, recalls the kernel $K_{1_a}$ to evaluate the Jacobian matrix $\mathbf{J}$ using the $\phi$-th parameterization and $\phi$-th set of initial concentrations as state of the system $\mathbf{X}$. As soon as $\mathbf{J}$ is calculated, thread $\phi$ applies Eq. 4 to estimate the dominant eigenvalue $\rho$ associated with the $\phi$-th parameterization and $\phi$-th set of initial concentrations;

- kernel $K_{1_a}$: each thread $j$, with $j = 0, \ldots, N-1$, calculates the values of the $j$-th row of $\mathbf{J}$ using the state of the system $\mathbf{X}$ and parameterization given as input.

Note that each thread $\phi$, with $\phi = 0, \ldots \Phi - 1$, exploits the DP to call the kernel $K_{1_a}$, by launching a novel grid of threads.

**Phase $\mathbf{P_3}$.** It implements the DOPRI5 method [2, 3, 4], an explicit Runge-Kutta integration algorithm of order 5 with variable step-size and stiffness control, used by the threads whose dominant eigenvalue $\rho$ is less than 500 to solve the system of ODEs. As a matter of fact, the $\Phi$ threads are partitioned in two different sets: $\mathcal{L}_{DOPRI}$ that contains the threads whose dominant eigenvalue $\rho$ is less than 500; $\mathcal{L}_{RADAU}$ that contains the remaining threads. If the DOPRI5 method fails in solving the system of ODEs characterized by a parameterization $\phi = 0, \ldots, \Phi$ and a set of initial concentrations, the $\phi$-th thread is moved into the set $\mathcal{L}_{RADAU}$. The DOPRI5 method relies on the following 12 kernels:

- kernel $K_2$: it is the main kernel implementing the DOPRI5 method. It is executed by each thread $d \in \mathcal{L}_{DOPRI}$, which uses its own parameterization as well as its own set of initial concentrations;
- kernel $K_{2_a}$: each thread $j$, with $j = 0, \ldots, N-1$, evaluates the $j$-th ODE using the state of the system $\mathbf{X}$ and parameterization given as input;
- kernel $K_{2_b}$: it exploits $N$ threads to update the vectors used to calculate the initial step-size of the DOPRI5 method;
- kernels $K_{2_c}$ – $K_{2_i}$: each kernel uses $N$ threads to update the vectors required by the DOPRI5 method to estimate the next step-size, using the Butcher tableau shown in Table 2 (left side);
- kernels $K_{2_j}$ and $K_{2_k}$: in each kernel, $N$ threads are launched to update the vectors involved in the spline approximation of the ODEs.

It is worth noting that each thread $d \in \mathcal{L}_{DOPRI}$ exploits the DP to call the kernels $K_{2_a}$ – $K_{2_k}$, by launching a novel grid of threads. Since data transfers between the CPU and the GPU are very time consuming, all the temporary results computed by FiCoS during this phase are stored on the global memory of the GPU.

**Phase $\mathbf{P_4}$.** This phase implements the RADAU5 method [5, 6], an implicit Runge-Kutta integration algorithm of order 5 with variable step-size. It is executed after the phase $P_3$, since the execution of some thread in that phase might fail. The RADAU5 method is based on the following 22 kernels:

- kernel $K_3$: it is the main kernel implementing the RADAU5 method. It is executed by each thread $r \in \mathcal{L}_{RADAU}$, which uses its own parameterization as well as its own set of initial concentrations;
- kernel $K_{1_a}$: each thread $j$, with $j = 0, \ldots, N-1$, calculates the values of the $j$-th row of $\mathbf{J}$ using the state of the system $\mathbf{X}$ and the parameterization given as input;
- kernel $K_{2_a}$: each thread $j$, with $j = 0, \ldots, N-1$, evaluates the $j$-th ODE using the state of the system $\mathbf{X}$ and parameterization given as input;
- kernel $K_{3_a}$: it uses $N$ threads to reinitialize the vectors involved during the Newton—Raphson method [9];
- kernels $K_{3_b}$ and $K_{3_c}$: each kernel takes advantage of $N$ threads to update the matrices—which are linearized—used during the linear system resolutions required by RADAU5 method;
- kernels $K_{3_d}$ and $K_{3_l}$: in each kernel, $N$ threads are launched to update the vectors used before and after the linear system resolutions required by RADAU5 method, relying on the Butcher tableau depicted in Table 2 (right side);

- kernels $K_{3_m}$ and $K_{3_o}$: these 3 kernels are used during the error estimation phase in which the new step-size is computed. Also in this case, each kernel takes advantage of $N$ threads to update the vectors required to calculate the new step-size based on the estimated error;
- kernels $K_{3_p}$ and $K_{3_s}$: in each kernel, $N$ threads are used to update the vectors involved in the spline approximation of the ODEs.

We highlight that both the matrix decompositions and the linear system resolutions were implemented exploiting the cuBLAS library [12]. Each thread $r \in \mathcal{L}_{RADAU}$ takes advantage of the DP to call the kernels $K_{1_a}$, $K_{2_a}$, $K_{3_a} - K_{3_s}$ launching a novel grid of threads. Since data transfers between the CPU and the GPU are very time consuming, all the temporary results computed by FiCoS during this phase are stored on the global memory of the GPU.

**Phase $\mathbf{P_5}$.** During this phase, for each simulation, the dynamics of the species to be sampled are written into output files. These output data (i.e., the concentration values of molecular species sampled at fixed time instants) are transferred to the host as soon as the phases $P_3$ and $P_4$ are completed.

### 4 Results

The results of the tests proposed in this section aim at showing the simulation accuracy of FiCoS, as well as the achieved speed-up, with respect to Livermore Solver of Ordinary Differential Equations (LSODA) [13], Variable-coefficient ODE (VODE) [14], cupSODA [15], and LASSIE (LArge-Scale SImulator) [16]. We leveraged the LSODA and VODE implementations provided by the SciPy scientific library (v.0.18.1) [17], and we used Python (v.2.7.12) and the NumPy library (v.1.11.2). cupSODA and LASSIE were downloaded from the github repositories `https://github.com/aresio/LASSIE` and `https://github.com/aresio/cupSODA`. All tests were performed on a workstation equipped with a CPU Intel Core i7-2600 CPU (clock 3.4 GHz) and 8 GB of RAM, running on Ubuntu 16.04 LTS. The GPU used in the tests was a Nvidia GeForce GTX Titan X (3072 cores, clock 1.075 GHz, RAM 12 GB), CUDA toolkit version 8 (driver 387.26).

#### 4.1 Simulation accuracy of FiCoS

The simulation accuracy of FiCoS was evaluated by using a CPU implementation of the numerical integrators ported on the GPU.

First, we compared the dynamics of the molecular species $EIF4EBP$ and $AMBRA$ of the RBM of the Autophagy/Translation switch based on the mutual inhibition of MTORC1 and ULK1[18], obtained by running LSODA, VODE, and FiCoS. The dynamics of this RBM was simulated by setting the initial value of $AMPK$ species to 90000, which allows for obtaining an oscillating behavior, thus making the system of ODEs stiff. We used the following settings for all siulators: $(i)$ absolute tolerance $\varepsilon_a = 10^{-12}$; $(ii)$ relative tolerance $\varepsilon_r = 10^{-6}$ and $(iii)$ maximum number of allowed steps equal to $10^4$. Fig. 4 shows that the dynamics obtained by FiCoS perfectly overlap those provided by LSODA and VODE, proving the correctness of the implementation of FiCoS as well as its accuracy.

Second, to assess the robustness of FiCoS with respect to the numerical integrators settings (i.e., absolute $\varepsilon_a$ and relative $\varepsilon_r$ tolerances), we run a batch of tests in which we changed one of these parameters at a time. The same tests were executed with LSODA and VODE, for a fair comparison. We evaluated the effect of these settings on both the quality of the output dynamics and the running time, considering the RBM of the Ras/cAMP/PKA signal transduction pathway in yeast [19, 20], which consists in 39 reactions and 33 molecular species, and it is characterized by an oscillatory behavior leading to the stiffness phenomenon.

The accuracy of the simulations outcome of LSODA, VODE, and FiCoS were assessed by comparing the dynamics of the molecular species cAMP simulated with different settings (see Table 3), against a reference dynamics obtained by running LSODA with the following settings: $\varepsilon_a = 10^{-12}$, $\varepsilon_r = 10^{-6}$ and maximum number of allowed steps equal to $10^4$. Figs. 5, 6, 7 show the results of this tests.

We observe that, in the case of FiCoS and VODE, all the dynamics are perfectly overlapped, as the tolerance values have no relevant effect on the quality of the output dynamics. Conversely, the dynamics obtained with LSODA changes with different tolerance settings. In the case of $\varepsilon_r = 10^{-2}$ and

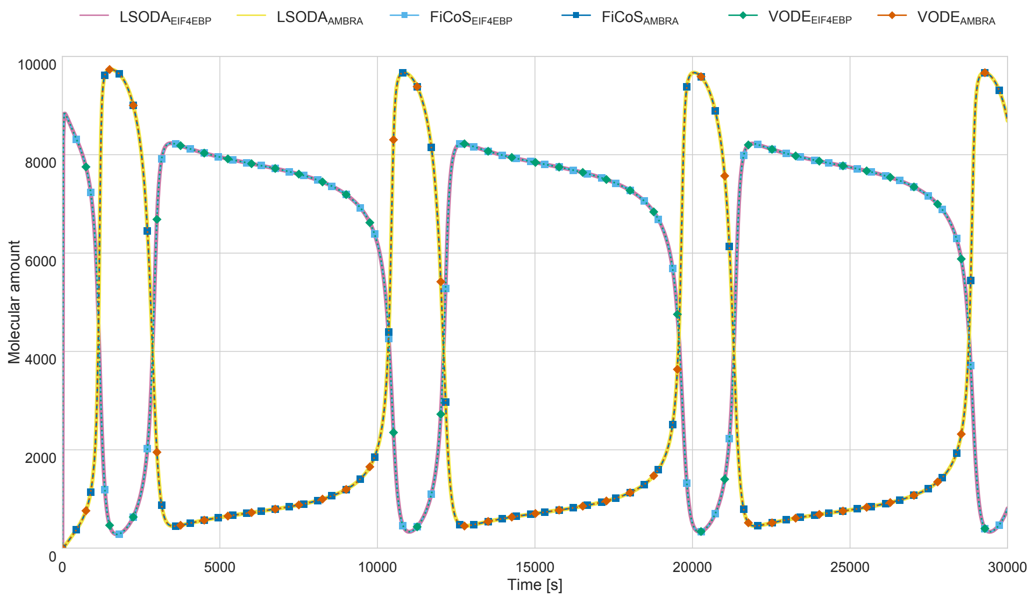


Figure 4: Comparison of the dynamics of the molecular species $EIF4EBP$ and $AMBRA$ of the model presented in [18], obtained by running FiCoS, LSODA and VODE, by using $\varepsilon_a = 10^{-12}$ and $\varepsilon_r = 10^{-6}$.

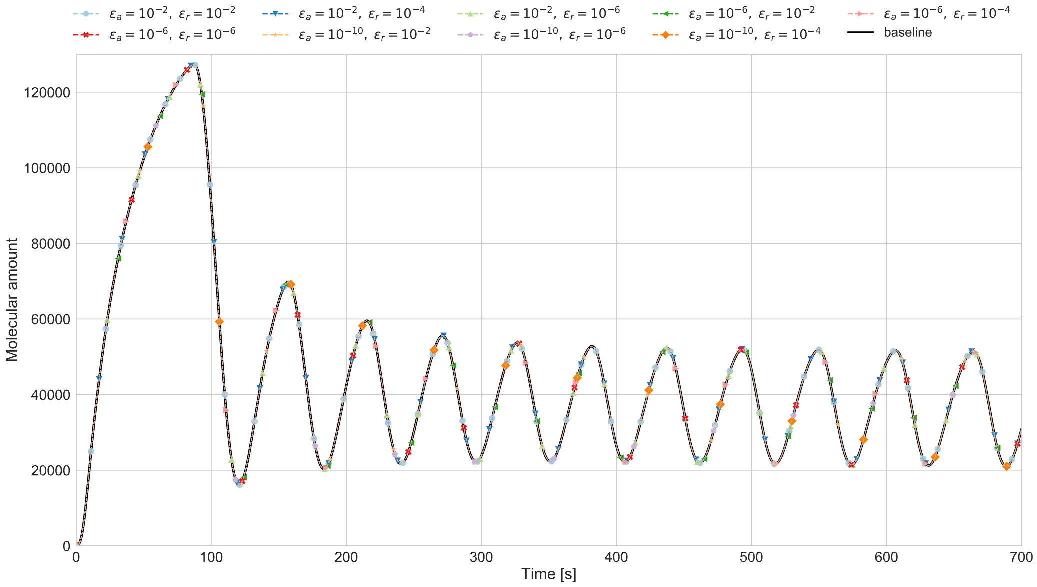


Figure 5: Comparison of the dynamic of the molecular species cAMP of the model of the RAS/cAMP/PKA signaling pathway in yeast, obtained by FiCoS varying $\varepsilon_a \in 10^{-2}, 10^{-6}, 10^{-10}$ and $\varepsilon_r \in 10^{-2}, 10^{-4}, 10^{-6}$.

$\varepsilon_a \in 10^{-2}, 10^{-6}$, the output dynamics do not perfectly overlap the reference dynamics (Fig. 7). This is probably caused by the automatically switching between the explicit and implicit families of integration methods. Since the explicit methods are not accurate for stiff systems, they are not able to properly integrate this ODE system, thus reducing the accuracy of the simulated dynamics.

Table 3 summarizes the running time required by LSODA, VODE and FiCoS to solve the systems of ODEs related to the RAS/cAMP/PKA signaling pathway in yeast, for each tested combination of $(\varepsilon_a)$ and $(\varepsilon_r)$ values. We observe that, while the running time of VODE remains almost constant with different tolerances values, FiCoS and LSODA performance can be dramatically affected by this choice,

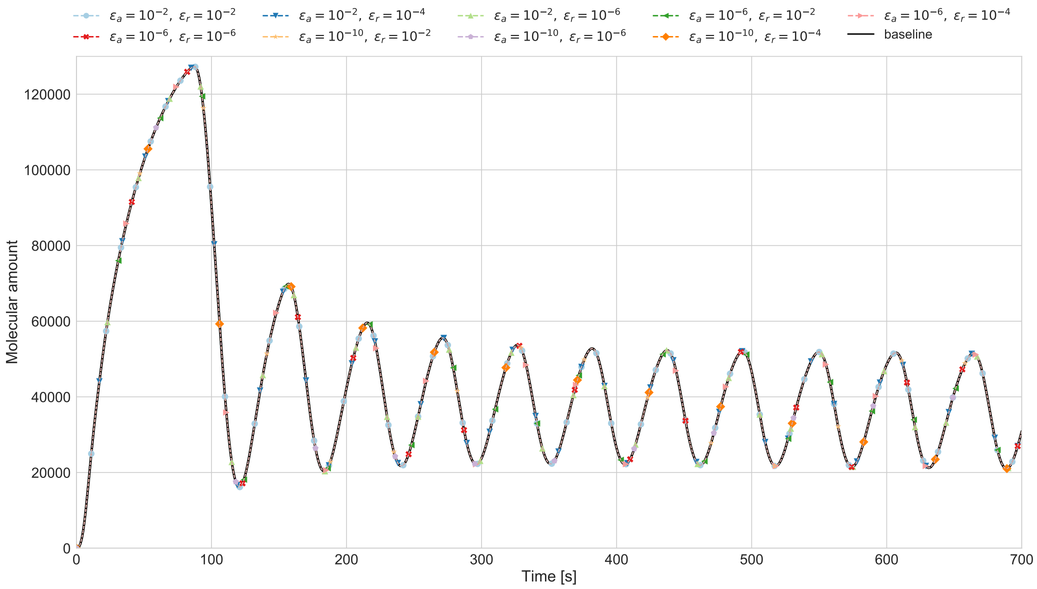


Figure 6: Comparison of the dynamic of the molecular species cAMP of the model of the RAS/cAMP/PKA signaling pathway in yeast, obtained by VODE varying $\varepsilon_a \in 10^{-2}, 10^{-6}, 10^{-10}$ and $\varepsilon_r \in 10^{-2}, 10^{-4}, 10^{-6}$.

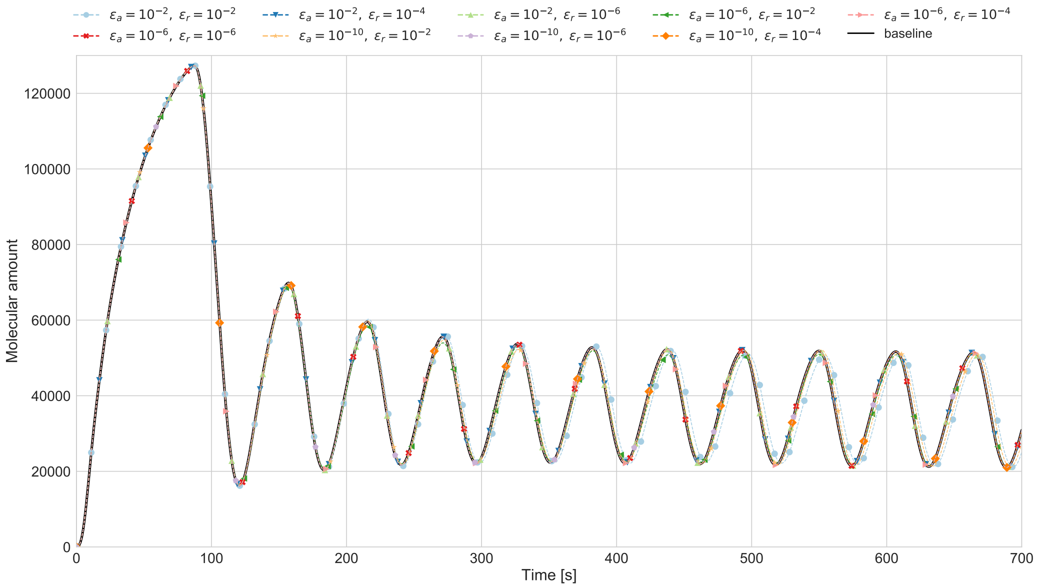


Figure 7: Comparison of the dynamic of the molecular species cAMP of the model of the RAS/cAMP/PKA signaling pathway in yeast, obtained by LSODA varying $\varepsilon_a \in 10^{-2}, 10^{-6}, 10^{-10}$ and $\varepsilon_r \in 10^{-2}, 10^{-4}, 10^{-6}$.

as the running times increase up to three times with some tolerance values combinations. To better investigate this aspect, we defined another batch of tests in which we varied the tolerances values and analyze the running time required to simulate the Autophagy/Translation switch based on the mutual inhibition of MTORC1 and ULK1 model, which is characterized by by 173 molecular species and 6581 reactions. Table 4 shows that the running time of VODE double from the combination $\varepsilon_a = \varepsilon_r = 10^{-2}$ to the combination $\varepsilon_a = 10^{-10}$ and $\varepsilon_a = 10^{-6}$, while the running time of LSODA and FiCoS triple. We also observe that the relative tolerance has a higher impact on the running time with respect to the

Table 3: Running time required by LSODA, VODE, and FiCoS to simulate the RBM of the RAS/cAMP/PKA signaling pathway in yeast, with different settings of absolute ($\varepsilon_a$) and relative ($\varepsilon_r$) tolerance.

| Settings | | Running time [s] | | |
|---|---|---|---|---|
| $\varepsilon_a$ | $\varepsilon_r$ | LSODA | VODE | FiCoS |
| $10^{-2}$ | $10^{-2}$ | 0.255 | 2.516 | 0.030 |
| $10^{-2}$ | $10^{-4}$ | 0.306 | 2.538 | 0.041 |
| $10^{-2}$ | $10^{-6}$ | 0.381 | 2.419 | 0.030 |
| $10^{-6}$ | $10^{-2}$ | 0.339 | 2.508 | 0.043 |
| $10^{-6}$ | $10^{-4}$ | 0.407 | 2.465 | 0.059 |
| $10^{-6}$ | $10^{-6}$ | 0.579 | 2.419 | 0.081 |
| $10^{-10}$ | $10^{-2}$ | 0.324 | 2.533 | 0.046 |
| $10^{-10}$ | $10^{-4}$ | 0.373 | 2.481 | 0.034 |
| $10^{-10}$ | $10^{-6}$ | 0.601 | 2.501 | 0.052 |

Table 4: Running time required by LSODA, VODE, and FiCoS to simulate the Autophagy/Translation switch based on the mutual inhibition of MTORC1 and ULK1 model, with different settings of absolute ($\varepsilon_a$) and relative ($\varepsilon_r$) tolerance.

| Settings | | Running time [s] | | |
|---|---|---|---|---|
| $\varepsilon_a$ | $\varepsilon_r$ | LSODA | VODE | FiCoS |
| $10^{-2}$ | $10^{-2}$ | 12.599 | 25.922 | 2.398 |
| $10^{-2}$ | $10^{-4}$ | 16.348 | 27.101 | 2.791 |
| $10^{-2}$ | $10^{-6}$ | 19.722 | 29.449 | 3.311 |
| $10^{-6}$ | $10^{-2}$ | 13.261 | 31.077 | 2.661 |
| $10^{-6}$ | $10^{-4}$ | 19.433 | 32.822 | 3.717 |
| $10^{-6}$ | $10^{-6}$ | 33.544 | 43.372 | 5.655 |
| $10^{-10}$ | $10^{-2}$ | 13.747 | 32.503 | 2.774 |
| $10^{-10}$ | $10^{-4}$ | 21.219 | 33.860 | 4.329 |
| $10^{-10}$ | $10^{-6}$ | 37.02 | 56.687 | 6.952 |

absolute tolerance: in the case of FiCoS, with $\varepsilon_a = 10^{-10}$, the running time triple from $\varepsilon_r = 10^{-2}$ to $\varepsilon_r = 10^{-6}$.

Finally, this tests show that the CPU implementation of FiCoS is faster than LSODA and VODE, with up to 84× speed-up, achieving in some cases more accurate dynamics.

#### 4.2 Computational performance

Starting from the results presented in the previous section, we present here the investigation of the computational performance of the GPU-powered implementation of FiCoS. The following simulations were executed with absolute tolerance $\varepsilon_a = 10^{-12}$, relative tolerance $\varepsilon_r = 10^{-6}$ and maximum number of allowed steps equal to $10^4$, which represent settings widely used in the literature as well as in well-known simulation software like COPASI [21]. We also highlight that in every simulation we save the dynamics of all species included in the RBMs. For each RBM, we performed an increasing number of simulations (up to 2048) and calculated both the integration time and the simulation time. The integration time indicates the running time spent by the numerical integration algorithms to solve the system of ODEs, while the simulation time is the overall running time required to perform a simulation, including the I/O operations (i.e., reading and writing operations). By using these data, we calculated the speed-up achieved by FiCoS with respect to the CPU-based ODE solver LSODA[13] and VODE[14], and the GPU-powered algorithms cupSODA [15] and LASSIE [16].

We carried out different batches of simulations using a set of "symmetric" synthetic RBMs of increasing size, ranging from 64 to 800 species and reactions, to evaluate the impact of the models size on the computational performance, and a set of "asymmetric" synthetic RBMs of increasing size, ranging from

21 species (reactions) and 64 reactions (species) to 267 species (reactions) and 800 reactions (species). The asymmetric RBMs were used to evaluate how much either the number of species or the number of reactions affect the performance of FiCoS.

**Symmetric models.** Figs. 8-11 present the results obtained with the batch of tests on symmetric RBMs. The panels in the figures report the speed-up values achieved by FiCoS, calculated considering the entire simulation time (a) and integration time only (b), with respect to LSODA, VODE, LASSIE, and cupSODA.

FiCoS provides a relevant reduction of the entire simulation time, achieving a 360× speed-up against LSODA (Fig. 8a), and a 487× speed-up against VODE (Fig. 9a) in the case of 2048 simulations of the $800 \times 800$ RBM. Note that the maximum speed-up value with respect to LSODA (366×) is reached in the case of 1024 simulations of the $800 \times 800$ RBM. We conjecture that for RBMs of larger size, the speed-up could be even higher with respect to a CPU-bound execution; however, we did not perform such tests due to the excessive memory requirements of LSODA and VODE implementations.

Since FiCoS was designed to exploit both a fine- and a coarse-grained strategy, the intra-GPU communication overhead—caused by the fine-grain kernels—makes the simulation to less efficient in the case of small size RBMs. Moreover, FiCoS performance can be hampered when a few simulations are executed. For instance, in the case of a single simulation of a RBM of size up to $256 \times 256$, FiCos is slower than both LSODA (Fig. 8a) and VODE (Fig. 9a). Anyway, while for a single simulation of small and medium RBMs a CPU-bound simulation should be preferred, in all other cases FiCoS represents the best solution.

Considering the performance achieved in the case of high numbers of simulations, we observed that an "excessive" parallelization might cause the saturation of the computing resources of the GPU. This is due to the computing resources required by the fine-grained kernels, exploited to parallelize the resolution of the system of ODEs, which increase along with the RBM size. As a matter of fact, the blocks of threads generated using the DP saturate the GPU resources, resulting in a decrease of the speed-up. In our tests, this phenomenon occurs for RBMs with more than 512 species and reactions, in the case of 2048 simulations.

Restricting the analysis of the performance of FiCoS to the integration time only, the portion of the running time spent to reading/writing operations of the input/output files, is excluded from the computation of the speed-up values. The speed-up achieved by the integration method implemented in FiCoS is smaller than the speed-up of the entire simulation time in the case of LSODA, with a speed-up up to 79×, while the speed-up with respect to VODE increases, up to 855× (see Figs. 8b and 9b, respectively). Overall, these results are consistent with the previous findings: FiCoS is faster than LSODA and VODE in all cases, except for the single simulation of the RBMs, and becomes less effective in the scarcity of GPU resources, e.g., when more than 2000 parallel simulations are executed.

Regarding the comparison of FiCoS against LASSIE, since the latter exploits only a single parallelization strategy, as shown in Fig. 10a, it is faster than FiCoS in the case of a single simulation (except for the RBM with size $512 \times 512$). Regarding the integration time, FiCoS always outperformed LASSIE, achieving a maximum speed-up equal to 5.57× (Fig. 10a). Considering tasks with multiple simulations, FiCoS resulted always faster than LASSIE; the highest speed-up is achieved in the case of 2048 simulations of the RBM with size $64 \times 64$, in which FiCoS is 298× faster than LASSIE for the simulation time (Fig. 10a), and 760× faster for the integration time (Fig. 10b).

We finally compared the performance of FiCoS against cupSODA, a GPU-powered simulator designed for small RBMs. cupSODA resulted faster with respect FiCoS in the case of a single simulation when less than 512 species and reactions are taken into account (see Fig. 11a). In the case of the smallest RBM considered in these tests (i.e., 64 species and reactions), FiCoS is more efficient only when more than 128 simulations are executed. In all remaining cases, FiCoS is the best choice, achieving a speed-up up to 7.35×. Regarding the integration time, FiCoS always outperformed cupSODA, reaching the maximum speed-up (17×) in the case of 128 simulations of the RBM with 800 species and reactions (see Fig. 11b). Note that, due to the required amount of memory, cupSODA failed to perform 2048 simulations of the RBMs with 640 and 800 species and reactions.

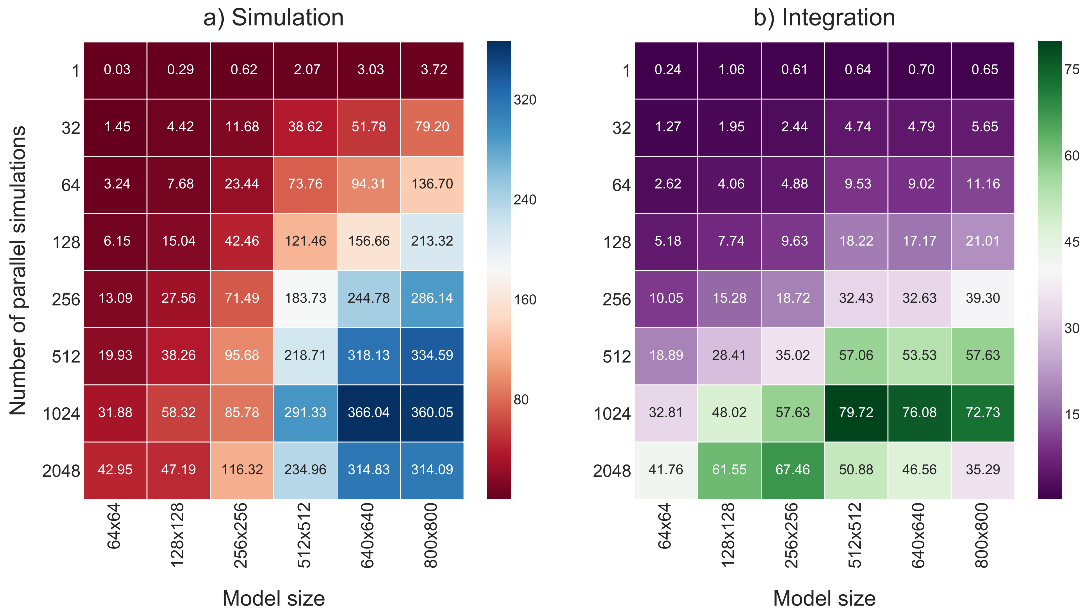


Figure 8: Speed-up provided by FiCoS with respect to LSODA[13] in the case of symmetric RBMs. The speed-up is analyzed by increasing both the size of the RBM (horizontal axis) and the number of parallel simulations (vertical axis). Panel a) and b) show the speed-up calculated considering the entire simulation time and numerical integration time, respectively.

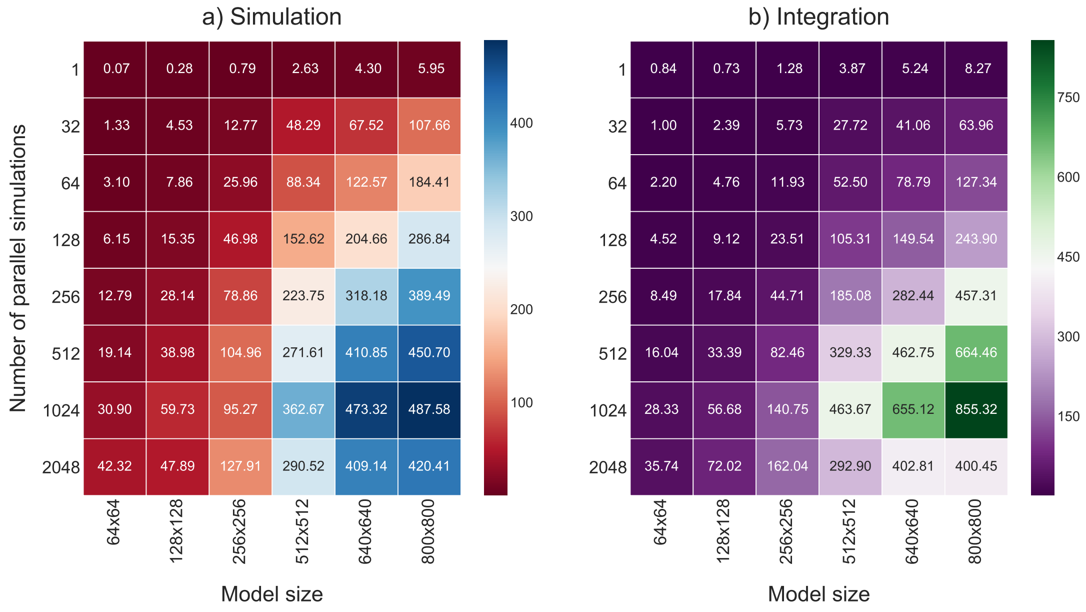


Figure 9: Speed-up provided by FiCoS with respect to VODE[14] in the case of symmetric RBMs. The speed-up is analyzed by increasing both the size of the RBM (horizontal axis) and the number of parallel simulations (vertical axis). Panel a) and b) show the speed-up calculated considering the entire simulation time and numerical integration time, respectively.

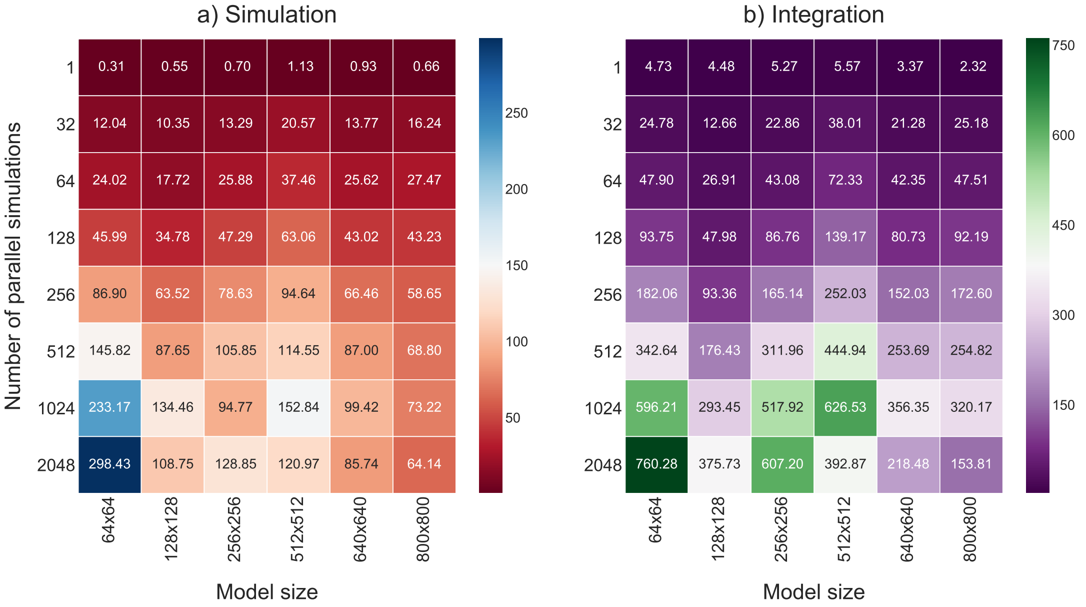


Figure 10: Speed-up provided by FiCoS with respect to LASSIE[16] in the case of symmetric RBMs. The speed-up is analyzed by increasing both the size of the RBM (horizontal axis) and the number of parallel simulations (vertical axis). Panel a) and b) show the speed-up calculated considering the entire simulation time the numerical integration time, respectively.

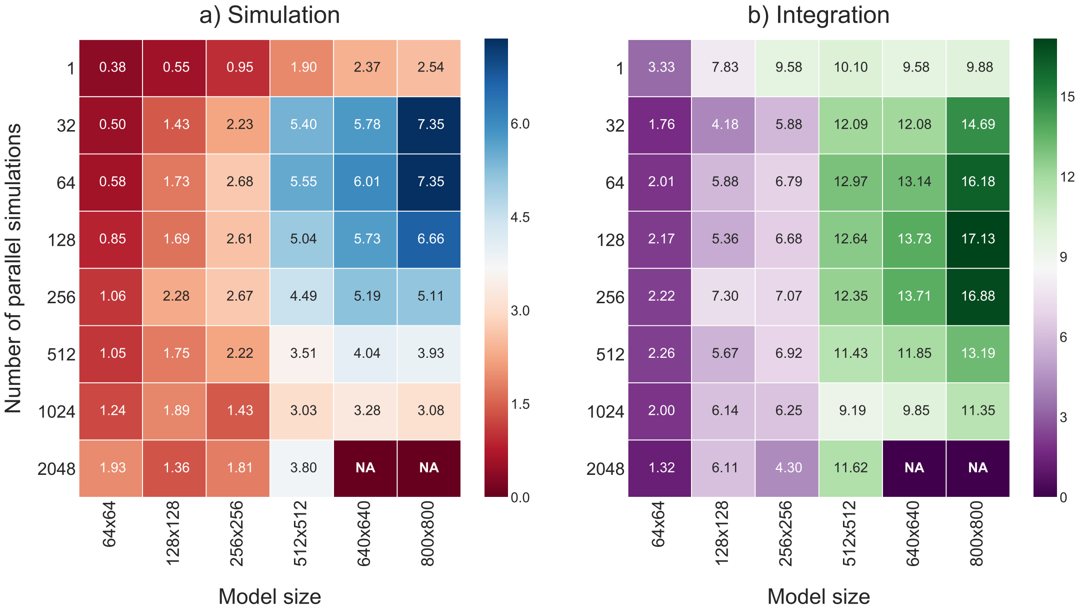


Figure 11: Speed-up provided by FiCoS with respect to cupSODA[15] in the case of symmetric RBMs. The speed-up is analyzed by increasing both the size of the RBM (horizontal axis) and the number of parallel simulations (vertical axis). Panel a) and b) show the speed-up calculated considering the entire simulation time and numerical integration time, respectively. NA values denote that cupSODA failed to perform the simulations due to the required amount of memory.

**Asymmetric models.** Figs. 12-19 present the results obtained with the batch of tests on asymmetric RBMs. The panels in these figures report the speed-up values achieved by FiCoS, calculated considering the entire simulation time (a) and the integration time only (b), with respect to LSODA, VODE, LASSIE, and cupSODA. In particular, to investigate how the number of species and reactions could affect the performance of the simulators (especially those exploiting the GPGPU computing), we generated a first set of asymmetric RBMs, in which the number of species is three times the number of reactions, and a second set having RBMs with opposite characteristics. These tests aim at analyzing the capabilities of FiCoS, which realizes both a fine- and coarse-grained parallelization strategy. Indeed, since each ODE corresponds to a different species, the higher the number of the species the higher the parallelization that can be obtained by exploiting a fine-grained strategy.

Conversely, since the number of reactions is roughly related to the length of each ODE, the number of operations that must be performed by each thread increases along with the number of reactions; therefore, the coarse-grained parallelization strategy might help mitigating the running time required.

Considering the first set of asymmetric RBMs, FiCoS was able to achieve speed-up values up to 413$\times$ and up to 508$\times$ with respect to LSODA and VODE, respectively, for what concerns the simulation time (Figs. 12a and 13a). Although these values are higher than those obtained in the case of symmetric RBMs (Figs. 8a and 9a), regarding the integration time, the speed-ups achieved by FiCoS are lower. LASSIE outperformed FiCoS only when a single simulation is performed. Considering the simulation time, the maximum speed-up is 255$\times$, while in the case of the integration time is 278$\times$ (Figs. 14 and 10). Finally, comparing FiCoS with cupSODA, the maximum speed-ups are halved with respect to case of symmetric RBMs (see Figs. 15 and 11). Moreover, cupSODA is faster than FiCoS for tasks involving up to 256 simulations of RBMs having size up to $256 \times 85$.

Considering the second set of asymmetric models, as shown in Figs. 16a and 17a, the speed-up of FiCoS is up to 252$\times$ and 277$\times$ with respect to LSODA and VODE, respectively, for the overall simulation time. Regarding the integration time, FiCoS achieved a speed-up up to 147$\times$ and up to 234$\times$ with respect to LSODA and VODE, respectively (Fig. 16b and Fig. 17b). Except for the single simulation of the RBM having size $21 \times 64$, LASSIE was always slower than FiCoS, the latter being capable of achieving speed-ups values up to 1100$\times$ considering the simulation time and 3666$\times$ considering the integration time (see Fig. 18). These tests evidenced the limitations of LASSIE, which implements only a fine-grained parallelization strategy, while FiCoS is capable of better exploiting the computing capabilities of the modern GPU. Finally, the comparison between FiCoS and cupSODA (Fig. 19) resulted in speed-up values up to 27$\times$ for the simulation time, and up to 60$\times$ for the integration time. cupSODA is more efficient than FiCoS only when simulating small-size RBMs; however, it failed to perform the simulations of RBMs of size $267 \times 800$ due to the required amount of memory.

Overall, these results confirmed that simulators exploiting only a fine- or a coarse-grained strategy are very inefficient in certain conditions, while the solution implemented in FiCoS allows for obtaining excellent performance in all computationally demanding tasks.

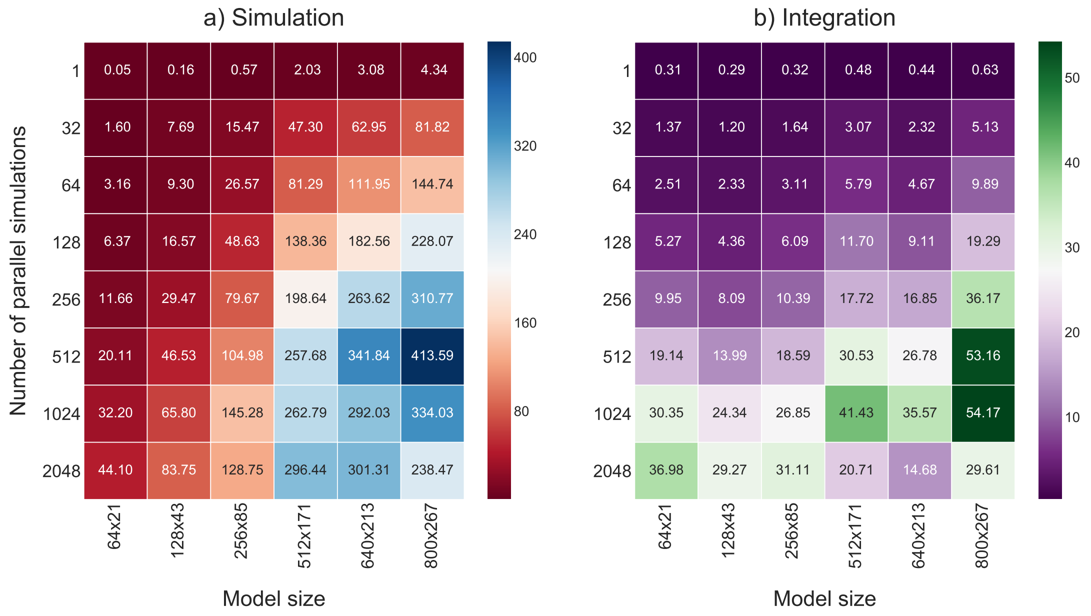


Figure 12: Speed-up provided by FiCoS with respect to LSODA[13] in the case of asymmetric RBMs. The speed-up is analyzed by increasing both the size of the RBM (horizontal axis) and number of parallel simulations (vertical axis). Panel a) and b) show the speed-up calculated considering the entire simulation time and numerical integration time, respectively.

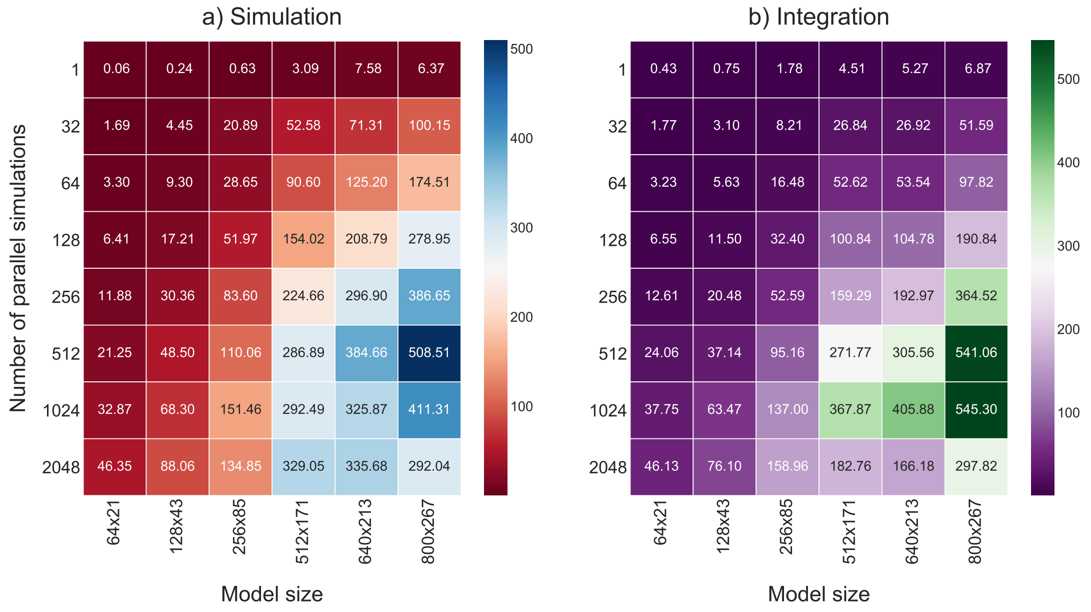


Figure 13: Speed-up provided by FiCoS with respect to VODE[14] in the case of asymmetric RBMs. The speed-up is analyzed by increasing both the size of the RBM (horizontal axis) and number of parallel simulations (vertical axis). Panel a) and b) show the speed-up calculated considering the entire simulation time and numerical integration time, respectively.

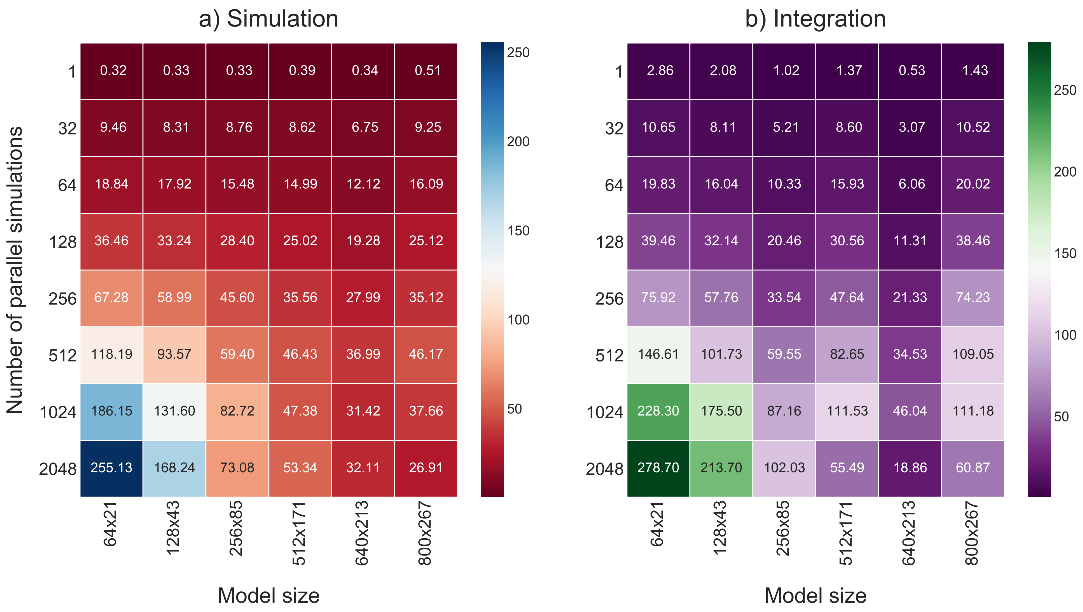


Figure 14: Speed-up provided by FiCoS with respect to LASSIE[16] in the case of asymmetric RBMs. The speed-up is analyzed by increasing both the size of the RBM (horizontal axis) and number of parallel simulations (vertical axis). Panel a) and b) show the speed-up calculated considering the entire simulation time and numerical integration time, respectively.

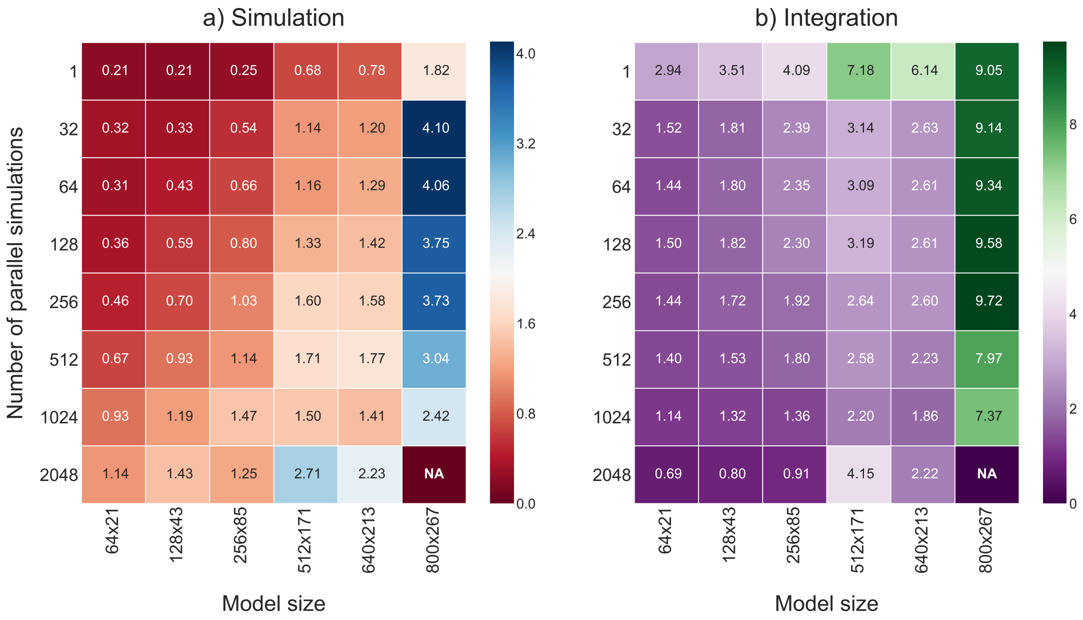


Figure 15: Speed-up provided by FiCoS with respect to cupSODA[15] in the case of asymmetric RBMs. The speed-up is analyzed by increasing both the size of the RBM (horizontal axis) and number of parallel simulations (vertical axis). Panel a) and b) show the speed-up calculated considering the entire simulation time and numerical integration time, respectively. NA values denote that cupSODA failed to perform the simulations due to the required amount of memory.

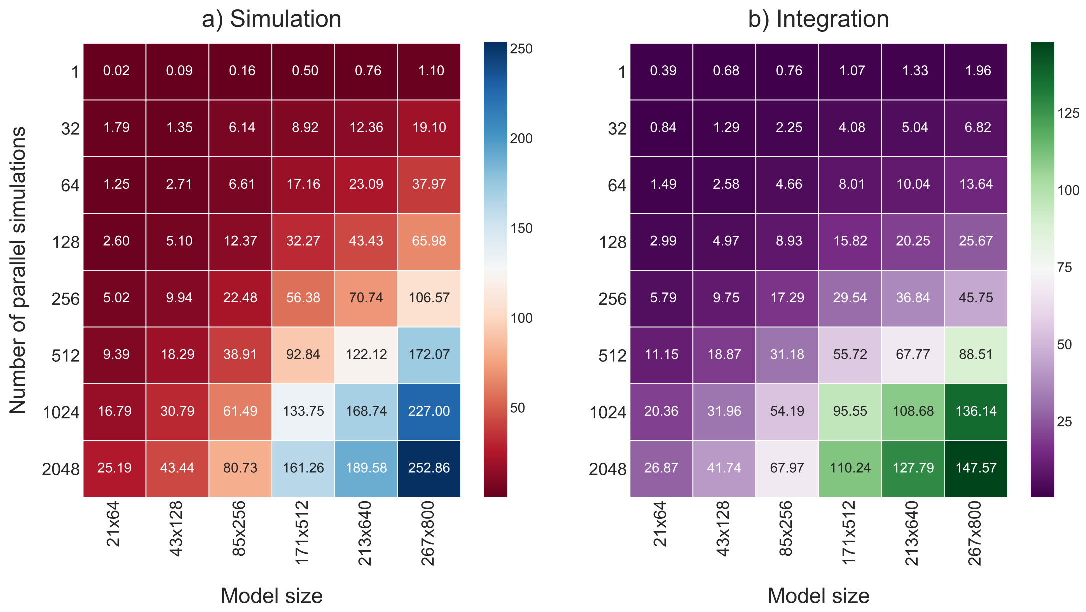


Figure 16: Speed-up provided by FiCoS with respect to LSODA[13] in the case of asymmetric RBMs. The speed-up is analyzed by increasing both the size of the RBM (horizontal axis) and number of parallel simulations (vertical axis). Panel a) and b) show the speed-up calculated considering the entire simulation time and numerical integration time, respectively.

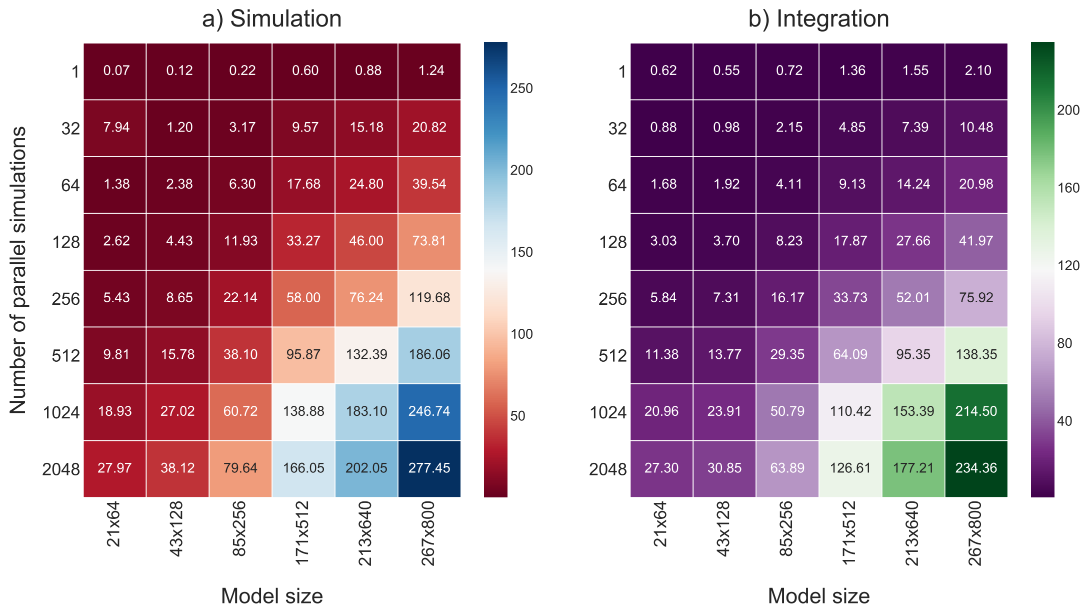


Figure 17: Speed-up provided by FiCoS with respect to VODE[14] in the case of asymmetric RBMs. The speed-up is analyzed by increasing both the size of the RBM (horizontal axis) and number of parallel simulations (vertical axis). Panel a) and b) show the speed-up calculated considering the entire simulation time and numerical integration time, respectively.

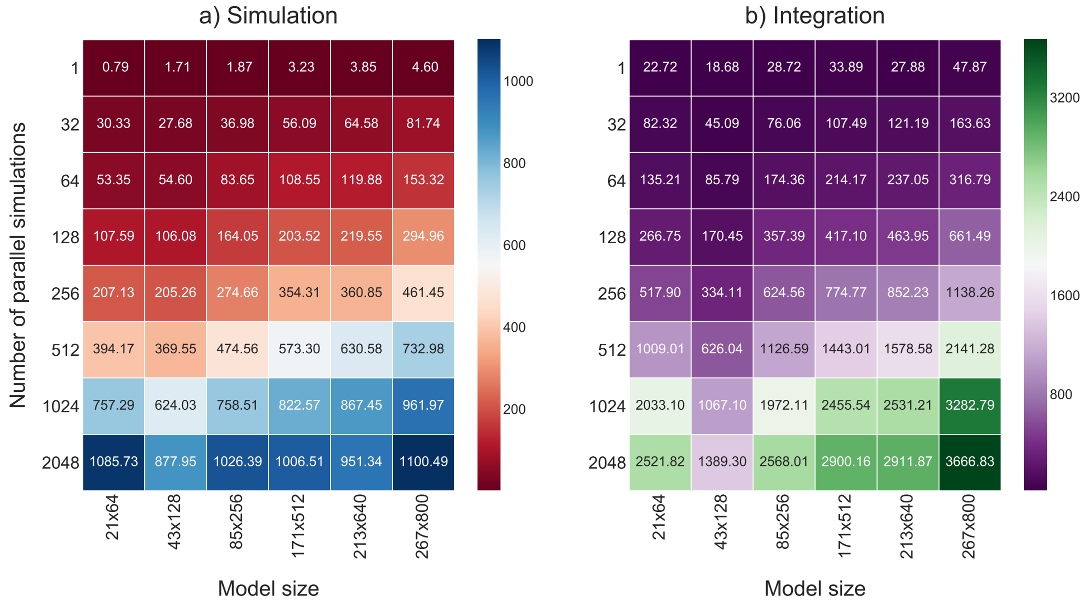


Figure 18: Speed-up provided by FiCoS with respect to LASSIE[16] in the case of asymmetric RBMs. The speed-up is analyzed by increasing both the size of the RBM (horizontal axis) and number of parallel simulations (vertical axis). Panel a) and b) show the speed-up calculated considering the entire simulation time and numerical integration time, respectively.

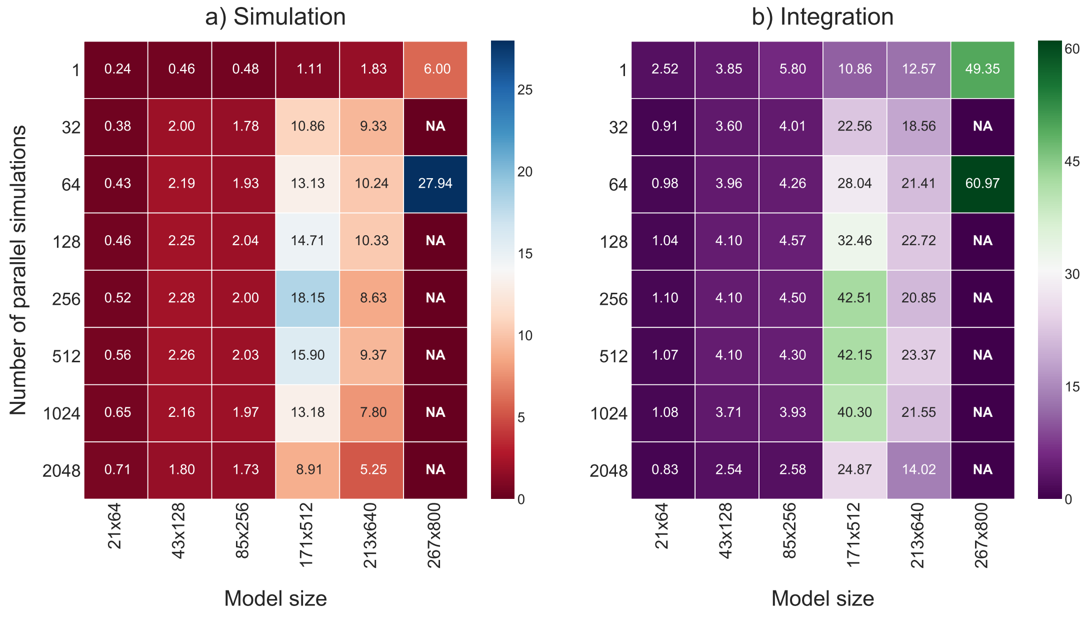


Figure 19: Speed-up provided by FiCoS with respect to cupSODA[15] in the case of asymmetric RBMs. The speed-up is analyzed by increasing both the size of the RBM (horizontal axis) and number of parallel simulations (vertical axis). Panel a) and b) show the speed-up calculated considering the entire simulation time and numerical integration time, respectively. NA values denote that cupSODA failed to perform the simulations due to the required amount of memory.

[11] Nvidia. CUDA C Programming Guide, version 8.0, 2016.

[12] Nvidia. cuBLAS library 8.0, 2015.
