## Supplementary Text S2 for "FiCoS: a fine-grained and coarse-grained GPU-powered deterministic simulator for biochemical networks"

### Input files

To correctly perform a simulation of a reaction-based model (RBM), FiCoS requires a list of input files, as reported in Table 1. For each file, we specify: (*i*) a brief description of the file content; (*ii*) the format of the file; (*iii*) a flag that indicates if the file is mandatory; (*iv*) the default values used by FiCoS, when applicable. Notice that all indices are 0-based and the files are in Tab-Separated Value (TSV) format. In the table, $N$ is the number of chemical species, $M$ the number of reactions involved in the model, and $E$ the number of required simulations. For each simulation, $T$ represents the number of sampling time instants in which the dynamics must be saved. FiCoS can accept a single parameterization consisting in the initial amounts of all species (either `M_0` or `MX_0`) and all kinetic constants (either `c_vector` or `c_matrix`), as well as multiple parameterizations described by a set of the initial amount of the species (`MX_0`) and a set of kinetic constants (`c_matrix`). Finally, FiCoS allows for computing in parallel a difference between the obtained simulations and a target series (`c=ts_matrix`). Specifically, FiCoS implements the fitness function described in [1].

Table 1: FiCoS input files

| *File name* | *Content* | *Format* | *Optional* | *Default* |
|---|---|---|---|---|
| `alphabet` | Vector containing the chemical species names | $N$ columns | Yes | $X_j$, with $j$=0, . . . , $N$-1 |
| `left_side` | Stoichiometric matrix of the reactants[1] | $M$ rows, $N$ columns | No | |
| `right_side` | Stoichiometric matrix of the products[2] | $M$ rows, $N$ columns | No | |
| `c_vector` | Vector of kinetic constants[3]. | $M$ rows | Yes, if `c_matrix` is provided | |
| `c_matrix` | Matrix of kinetic parameters[3] | $E$ rows, $M$ columns | Yes, if `c_vector` is provided | |
| `M_0` | Vector of the initial amounts of chemical species[4] | $N$ columns | Yes, if `MX_0` is provided | |
| `MX_0` | Matrix of the initial amounts of chemical species[4] | $E$ rows, $N$ columns | Yes, if `M_0` is provided | |
| `t_vector` | Vector of sampling time instants | $T$ rows | No | |
| `M_feed` | Matrix of the chemical species whose values must be kept constant[5] | $N$ columns or<br>$E$ rows, $N$ columns | Yes | $\mathbf{0}$ |
| `modelkind` | Type of input model[6] | {stochastic, deterministic} | Yes | deterministic |
| `volume` | Reaction volume of the system[7] | real number | Yes | |
| `cs_vector` | Vector of chemical species to be saved | $K \leq N$ rows | Yes | All chemical species |
| `atol_vector` | Vector of absolute error tolerances | $N$ rows | Yes | $\mathbf{10^{-12}}$ |
| `rtol` | Relative tolerance | real number | Yes | $10^{-6}$ |
| `max_steps` | Maximum number of allowed integration steps | real number | Yes | 10000 |
| `ts_matrix` | Matrix of target time series. The first column must be equal to `t_vector`. | $T$ rows, $K+1$ columns (with $K \leq N$) | Yes | |

[1]Left-hand side of the reactions.
[2]Right-hand side of the reactions.
[3]Both deterministic or stochastic kinetic constant values, associated with the reactions, are allowed (see `modelkind` file).
[4]Both concentration (real numbers) or molecular amount (integer number) values are allowed (see `modelkind` file).
[5]This vector/matrix indicates the chemical species whose amounts must be kept constant throughout the simulation (i.e., they are assumed to be constantly fed into the system). The values in this vector/matrix are equal to 0 if the species can vary in time, they are equal to 1 otherwise (keeping the values equal to the related values in `M_0` or `MX_0`).
[6]Deterministic: amount of chemical species given as concentration values, reaction parameters specified as deterministic reaction rates. Stochastic: amount of chemical species given as integer numbers of molecules, reaction parameters specified as stochastic constants.
[7]Required if the model is defined as stochastic in the `modelkind` file. It is exploited to convert the stochastic kinetic constants into the deterministic formulation as well as the molecular amounts (integer number) into concentrations (real numbers).
